# Inhibition of Xanthine oxidase by 1-*O*-methyl chrysophanol, a hydroxyanthraquinone isolated from *Amycolatopsis thermoflava* ICTA 103

**DOI:** 10.1101/2023.03.04.531071

**Authors:** Uma Rajeswari Batchu, Bharati Reddi, Joshna Rani Surapaneni, Prakasham Reddy Shetty, Sunil Misra, Anthony Addlagatta

## Abstract

Hyperuricemia caused by elevated levels of serum uric acid is responsible for implication of gout and other associated disorders that influence the human health. So far, Xanthine oxidase (XO) inhibitors are the choice of first line drugs for the treatment of hyperuricemia. The objective of the present study was to isolate a potent XO inhibitor from the actinobacteria and to evaluate its inhibitory mechanism. Initially, XO was isolated from bovine milk using standard protocol and enzyme kinetics were evaluated. Thereafter, culture filtrates of actinobacteria (*Amycolatopsis thermoflava* ICTA 103), *Streptomyces luteireticuli* ICTA 16, *Streptomyces kurssanovii* ICTA165 and *Amycolatopsis lurida* ICTA 194) were screened for XO inhibition using *in vitro* qualitative NBT plate assay followed by extraction and purification of potent inhibitor 1-*O*-methyl chrysophanol (OMC), from the culture filtrate of *Amycolatopsis thermoflava* ICTA 103, which belongs to hydroxy anthraquinones (HAQ) family. Further, *in silico* molecular model building was performed to study the binding affinity of OMC towards XO followed by quantitative *in vitro* spectroscopic assays. The molecular building study explored the mechanistic view of binding interaction between inhibitor & enzyme and the results were corroborates with the *in vitro* kinetic study. The *in vitro* results revealed the significant enzyme inhibition potential of OMC with an IC_50_ and *K*_*i*_ value of 24.8 ± 0.072 µM & 2.218 ± 0.3068 µM respectively. These results are comparable to standard allopurinol, however, more significant than its structural analog, chrysophanol. The kinetic analysis revealed that OMC is a reversible slow binding inhibitor and the Lineweaver - Burkplot analysis showed mixed type inhibition of OMC against XO. These results are in agreement with chrysophanol. Findings of this study proposed a new derivative of HAQ in the pipeline of hyperuricemia therapeutic drug candidates.

## Introduction

Hyperuricemia is considered as a foremost cause of gout and associated with other complex co morbidities such as metabolic as well as kidney and cardiovascular disorders which further lead to an increased risk of mortality ^[1]^. It has been described as a clinical condition associated with elevated plasma uric acid levels higher than 6.8 mg/dl at physiological temperature (37 °C) and pH 7.4 caused by overproduction or under excretion of serum uric acid. Uric acid is the end product of purine catabolism and is excreted through urine in humans and higher primates under normal or healthy condition ^[2]^. Xanthine oxidase (XO) is the key enzyme that converts hypoxanthine to xanthine and ultimately to uric acid along with reactive oxygen species/reactive nitrogen species (ROS/RNS) as byproducts of oxidative hydroxylation ^[3]^. However, under pathological conditions, the irreversibly accumulated uric acid causes hyperuricemia which predisposes into gout, whereas the ROS/RNS generated as by products may lead to inflammation, atherosclerosis, myocardial infarction and cancer ^[4-5]^. Therefore, selective inhibition of XO will have a promising role in the treatment of gout and other XO associated diseases. The purine analog, allopurinol and non-purine thiazole derivatives, febuxostat ^[6]^ and topiroxostat ^[7]^ are commonly used drugs as XO inhibitors for many years and showed constructive effects in the treatment of gout and other XO associated diseases ^[8-10]^. However, the frightful adverse effects such as Steven-Johnson syndrome and toxic epidermal necrolysis associated with these medication necessitates the search for novel natural XO inhibitors with potent activity and minimal toxicity ^[11-12]^.

Researchers worldwide evaluated various biologically originated novel compounds as XO inhibitors and noticed that few hydroxyl anthraquinones (HAQ) and their derivatives viz., anthrarobin, anthragallol, aloe-emodin, and chrysophanol have been explored as potent XO inhibitors. Over the above, earlier literature reports explored the potent anti-inflammatory potential of HAQ along with other bio functionalities and observed to be promising with minimal side effects ^[13-15]^. Maksimovic et al., 2010 reported that compounds having antioxidant and anti-inflammatory potential are also competent to treat gout and associated inflammatory diseases owing to their ability to reduce uric acid & ROS levels ^[16]^. Therefore, the authors streamlined that 1-*O*-methyl chrysophanol (OMC), an HAQ isolated from actinobacteria, *Amycolotopsis thermoflava* ICTA 103 may also inhibit the XO enzyme activity owing to its reported antioxidantas well as anti-inflammatory potential ^[17]^. Therefore, in the present study, OMC was extracted and purified from the culture filtrates of *A. thermoflava* ICTA 103 and studied for its inhibition potential against XO, isolated from the bovine milk. In addition, the biochemical mechanism of inhibition was evaluated using kinetic and Lineweaver-Burkplot analysis. *In silico* molecular model building was performed to reveal the binding efficiency.

## Materials and Methods

### Chemicals and reagents

The chemicals used for enzyme assays (allopurinol, xanthine, chrysophanol, tris-HCl and chloroform, methanol and ethylenediaminetetraacetic acid (EDTA), calcium phosphate gel (Type II: Neutral Brushite) were procured from Sigma Aldrich (St Louis Mo, USA). Sodium salicylate, Dithioerythritol (DTE) were purchased from Hi Media laboratories (Chennai, India). All chemicals and reagents used were of analytical grade.

### Culture collection and maintenance

The glycerol stocks of *Amycolatopsis thermoflava* ICTA 103, *Streptomyces luteireticuli* ICTA 16, *Streptomyces kurssanovii* ICTA165 and *Amycolatopsis lurida* ICTA194 from IICT were revived using nutrient broth to be used readily. The strains were further maintained on Actinomycetes Isolation Agar (AIA) plates at 4° C for future studies.

### Isolation and partial purification of XO from bovine milk

The XO enzyme was isolated from the 1 liter bovine milk by using the modified method of Nakamura &Yamazaki ^[18]^. In brief, fresh bovine milk was purchased from the local dairy form (Vijaya Dairy, Hyderabad). The procured milk was incubated with 2.5 mM DTE, 1 mM EDTA and 1.25 mM sodium salicylate by stirring followed by cooling at 4°C. The cooled mixture was subjected for centrifugation at 3000 × g for 20 min. To the upper creamy layer, an equal volume of 0.2 M potassium phosphate buffer, containing 2.5 mM DTE, 1 mM EDTA and 1.25 mM sodium salicylate was added and incubated at 4°C. To this cooled mixture, cold (−20°C) butanol was slowly added to give 15% (v/v) concentration while stirring. Finally, 15% (w/v) solid ammonium sulphate ((NH_4_)_2_SO_4_) was supplemented and stirred constantly and incubated for 1 h followed by centrifugation at 13 000 × g for 20 min. The aqueous lower phase was separated and stirred again with 15% (w/v) solid (NH_4_)_2_SO_4_. The resultant suspension was allowed to stand at 4°C for 2 h and the top phase was collected and centrifuged at 10000 × g for 30 min. Subsequently the collected top phase was suspended in an estimated smaller volume of 0.2 M potassium phosphate buffer (pH 6) containing 2.5 mM DTE, 1 mM EDTA and 1.25 mM sodium salicylate and dialyzed against the similar buffer overnight under continuous stirring at 4°C.Theprecipitate formed during dialysis was removed by centrifugation at 20,000 × g for 1 h, producing a supernatant of ‘crude enzyme’.

The crude enzyme was applied to a column (1.5 cm × 15 cm) packed with calcium phosphate gel (Type II: Neutral Brushite) which was previously equilibrated in buffer A. The column was washed with this buffer until no more protein was eluted. Bound XO was then eluted from the column by using Buffer A, containing, additionally, 5% (w/v) (NH_4_)_2_SO_4_ followed by dialysis of combined protein fractions overnight against suitable buffer. The protein fraction was analyzed for purity using 12% SDS-PAGE gel electrophoresis method. The concentration of the enzyme was determined by using Nano drop (Nano drop 1000 Spectrophotometer, Thermo scientific, U.S.A). Further, one-milliliter enzyme aliquots were prepared and preserved at -80° C until further studies. Thereafter, the activity of enzyme was calculated by spectrophotometric assay at 295nm using xanthine as a substrate before each analysis.

### Optimization of Enzyme kinetics

#### Effect of pH

The effect of pH on the XO activity was determined using different buffers having various pH conditions: acetate buffer (pH 4.5, 5.0, 5.5), sodium phosphate buffer (pH 6, 6.5,7.0), potassium phosphate buffer (7.5) tris (pH 8.0, 8.5) and sodium carbonate (pH 9.0, 9.5) The reaction was executed using 50mM buffer with 0.03 µM enzyme and 500 µM xanthine as a substrate in the 96-well flat bottom micro titer plate (Tarsons, India) using a multimode plate reader (TECAN, Austria) at 295 nm.

### Effect of metal ions on XO activity

Being a metalloenzyme, the hydroxylation of purines occur at the M_O_ cofactor in the C-terminal domain of XO. A small amount of the isolated bovine milk XO exists in inactive demolybdo form, however, the percentage depends on the nutritional status of the cows ^[19]^. Therefore, the present study was focused to test the metal selectivity towards enzyme activity in 50mM potassium phosphate buffer (pH 7.5) using various divalent metal cations.

### Enzyme kinetics

The enzymatic assay for XO was performed as per the modified method of Cos et al. using above optimized conditions ^[20]^. Briefly, 100 µl of optimized enzyme reaction mixture was prepared containing 50 mM potassium phosphate buffer (pH 7.5), 0.03 µM XO and the reaction was started by adding 500 µM xanthine as a substrate. The absorbance was monitored for 30 min at 295nm using multimode reader (Tecan, Austria).

In the supporting experiment, K_M_ and Vmax values of XO were determined at varied concentrations of xanthine as a substrate (3 to 1000 µM). The result was calculated using GraphPad prism (GraphPad Software 8.0, San Diego CA) by applying non-linear curve fit analysis (figure 1).

**Figure 1:**
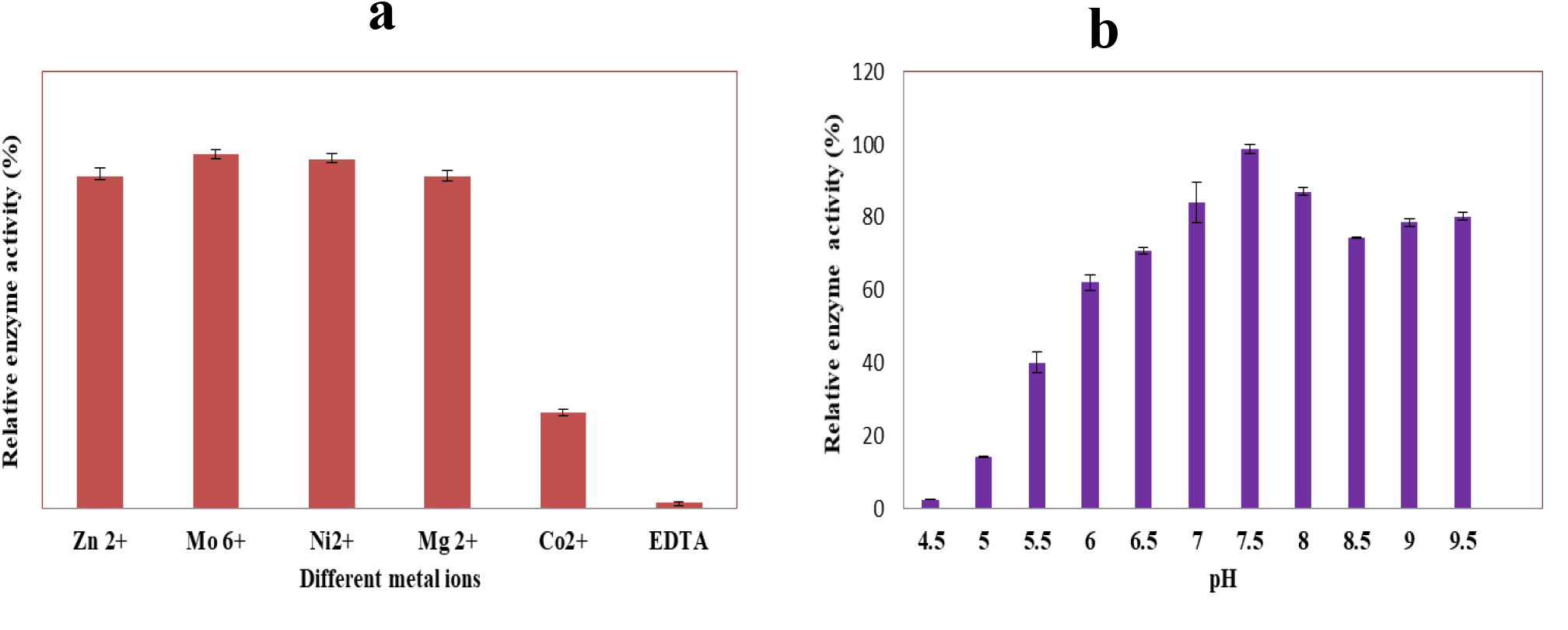
Enzyme kinetics optimization a) Study of metal ion effect on XO activity b) pH effect on XO activity.

Whereas the turnover number and Kcat values were determined from the formula

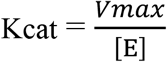

Vmax =maximum velocity and [E]= Enzyme concentration. The data was shaped into Michaelis-Menten equation (*V* = Vmax [S] / Km + [S])

### Fermentative production of culture filtrates of actinobacteria

Different species of actinobacteria, *S. luteireticuli* ICTA 16, *S. kurssanovii* ICTA165, *A. thermoflava* ICTA 103 and *A. lurida* ICTA194 were subjected for fermentation at 30 °C (*S. luteireticuli* ICTA 16, *S. kurssanovii* ICTA165 and *A. lurida*) at 40 °C (*A. thermoflava* ICTA103) and incubated for 5 days. After incubation, theculture broth was centrifuged (REMI C-24BL, India) at 10,000×g to collect supernatant which was further subjected for qualitative screening of XO inhibition by plate assay.

### Qualitative screening for XO inhibition (NBT plate assay)

The culture filtrates of selected strains were screened for XO inhibition by modified qualitative Xanthine-Nitroblue tetrazolium chloride (NBT) plate assay method ^[21]^. The Xanthine-NBT plates were prepared using 1.5% agar, 1.5 mg/ml Xanthine, 0.11 mg/ml NBT in potassium phosphate buffer (pH-7.5). A 5 mm wells were prepared aseptically with sterile stainless steel borer. Subsequently, 20µl of reaction mixture consisting of 10µl of test compound and 10 µl of 0.04 U of XO prepared in 50 mM potassium phosphate buffer (pH 7.5) was dispensed into each well and incubated overnight at 37^°^ C. The control well was made to occupy with DMSO and 0.04 U of enzyme. Allopurinol (2 mM) was used as a positive control. The appearance of blue color halo zone in control well indicates the XO activity whereas reduction in diameter of blue color halo zone in test and positive control wells in comparison to control indicates XO inhibition. All the tests were carried out in triplicates. The diameter of the zone was recorded and values were represented as mean ±SD values.

### Isolation and purification of OMC from *Amycolatopsis thermoflava* ICTA 103

Owing to the significant XO inhibition potential of the *Amycolatopsis thermoflava* ICTA 103culture filtrate, further studies were performed for isolation and purification of metabolite from the culture filtrate of the strain. The procedure for isolation and purification was described elsewhere ^[17]^. Briefly, the culture supernatant was treated with resin to adsorb the compound followed by resin washing and drying. The compound was extracted from dried resin at room temperature by using methanol. The crude extract was dried using rotavac and subjected for purification using silica gel column chromatography (100-200 mesh size) with a methanol-chloroform system (5:95 v/v). Further, the purified compound was characterized by using spectroscopic analysis including NMR, FT-IR, HR-MS and subjected to further quantitative XO inhibition studies (Figure S2-S6).

### Molecular model building

The molecular model building studies of OMC and chrysophanol were performed at the quercetin binding site of XO and comparisons were drawn using coot model-building tool ^[22]^. The 3D Structures of ligands were prepared using Chemdraw software and the crystal structure of bovine milk XO (3NVY.pdb) was downloaded from the Protein Data Bank (PDB).

### Quantitative method

#### Xanthine-XO scavenging assay

The assay was conducted according to modified method of Kapoor and Saxena ^[23]^. Initially, 20 µl of XO (0.04 U) enzyme solution was pre incubated with 20 µl test or standard compounds at 37^°^ C for 1 hr. Thereafter, the reaction was initiated by adding the 30 µl of xanthine (6.7 mM) followed by 30 µl of NBT (0.25mM) and further incubated for 30 min. The amount of the formazanformed was estimated by measuring the absorbance at 575 nm using a multimode plate reader (TECAN, Austria). Allopurinol and chrysophanol were used as pharmacological and structural standards, respectively. The control reaction mixture consists of xanthine, XO and NBT (without inhibitor). All the reactions were performed in triplicates and the results were expressed as mean ± SD.

### *In vitro* XO inhibition assay (Uric acid estimation assay)

The spectrophotometric method of XO inhibition was performed as per the modified method of Cos et al., described in the enzyme kinetics ^[23]^. Initially, the 100 µl of reaction mixture consisted of inhibitors (100 µM), 50mM potassium phosphate buffer pH 7.5, 0.03 µM enzyme and incubated for 15 min at 25^°^C. Later on, the reaction was initiated by adding 500 µM of xanthine as a substrate and the whole reaction mixture was incubated at 25^°^ C for 30 min. Thereafter, the reaction was terminated by adding 1N HCl following the estimation of the concentration of uric acid by measuring its absorbance at 290 nm using UV-Spectrophotometer (TECAN, Austria). Allopurinoland chrysophanolwere used as a pharmacological and structural standards respectively. The percentage inhibition of XO was calculated by following formula:

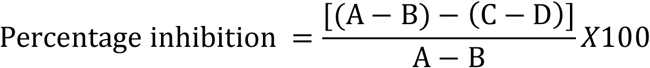

Where A is the OD at 290 nm with enzyme but without sample, B is the OD at 290 nm without sample and enzyme, C is the OD at 290 nm with sample and enzyme, and D is the OD at 290 nm with sample but without enzyme.

### Inhibition kinetics

Stock solutions of inhibitors (allopurinol, chrysophanol and OMC) were prepared using dimethyl sulfoxide (DMSO). Initially, the percentage XO inhibition of all the inhibitors was evaluated at 100µM concentration. Consequently, the inhibition constant *Ki* and IC_50_ were determined. The *K*_*i*_ value was calculated using Morrison method (at varied concentration of inhibitor) by fitting the data with non-linear regression using the formula below ^[24]^.

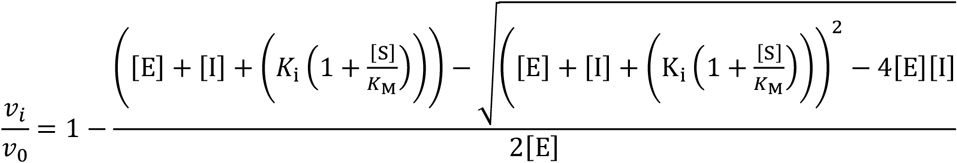

### Study of inhibition mode

#### Time dependent assays

The study was performed as per the modified method described by Benitez et al. 2022 ^[25]^. The inhibition studies were performed with the inhibitors (allopurinol, chrysophanol and OMC) at two different assay conditions. In the first reaction, 50µM OMC, 400 µM chrysophanol and 0.15 µM allopurinol (10mM stocks of inhibitors were prepared with 20% DMSO) were pre-incubated with the XO (0.03 µM) for 15 min at 25°C. Whereas, the second reaction proceeds under the same conditions without enzyme-inhibitor pre-incubation. The control reactions omitted inhibitors were run in parallel. Next, the reactions were initiated by addition of xanthine and incubated for 30 min at 25°C. Further, reactions were stopped by 1N HCL to measure the uric acid formation at A_294_ using multimode reader (Tecan, Austria). All the experiments were performed in triplicates and the data was represented as mean ± SD.

### Irreversible inhibition assays (Covalent inhibition)

Briefly, all the inhibitors (OMC, chrysophanol and allopurinol) were pre-incubated with the enzyme(0.03 µM) for 30 min at 25°C using the similar concentrations of the time dependent assay (10mM stocks of inhibitors were prepared with 20% DMSO). Thereafter, the excess concentration of the inhibitors was removed by using dialysis against the same buffer. Next, the samples were diluted with the reaction buffer (1:1) and incubated with xanthine at 25°C for 30 min. The reactions were stopped by 1N HCL to measure the uric acid formation at A_294_ using multimode reader (Tecan, Austria). The assays were replicated thrice and the % of enzyme inhibition was calculated. **Lineweaver-Burk plots**

To study the non-covalent mode of enzyme inhibition, Lineweaver-Burk plots were constructed by varying the concentration of xanthine substrate (3 to 1000 µM) in the presence and absence of the inhibitor at the concentrations of IC_25 &_ IC_50_ ^[26-27]^. The inhibition mode was compared with the standards allopurinol and chrysophanol.The plots were designed between reciprocal of velocity on Y-axis and reciprocal of substrate concentration on X-axis.

## Results and Discussion

Owing to the importance of XO inhibition as a therapeutic approach for treating the hyperuricemia, authors used XO as a target molecule to isolate the enzyme inhibitors from the microbial isolates. Considering the bovine milk as one of the richest source of XO, the same was isolated ^[28]^ and used for identification of actinobacterial sources having potential to produce XO inhibitors and the one of the potent actinobacteria, *Amycolatopsis thermoflava* ICTA103 has been further studied for isolation and purification of XO inhibitor and evaluated for its structural properties.

### Isolation of XO from bovine milk and activity measurements

The XO was isolated from the bovine milk by using ammonium sulphate precipitation method (Figure S1) and purified with calcium phosphate gel column. The purified fractions has shown a two major bands at molecular mass of 140 & 135 kDa in SDS-PAGE gel which corresponds to 98% of the protein. The other faint bands represents the traces of impurities obtained by degradation of enzyme due to internal proteases (The concentration of the enzyme was determined as 16mg/ml. Thereafter, the enzyme was evaluated for its activity using xanthine as substrate and activity was observed as 0.7 U/mg protein.One unit of enzyme activity was defined as the amount of enzyme that converts one µmol of xanthine to uric acid per min under defined conditions. Similar results have been noticed by Ozer et al., where the activity of the isolated enzyme was reported as 1.086 U/mg protein which is slightly differed to the present study ^[29]^. This might be attributed to the source of milk and extraction conditions etc.

### Enzyme kinetics

XO is a molybdenum (M_O_^+6^) incorporated metalloenzyme which plays a key role in catabolism of purine nucleotides. Moreover, MO^+6^as a constituent of purified enzyme from bovine milk was well studied (Green & Beinert 1953). Although it was also reported that 5-26% of enzyme in bovine milk exists in inactive demolybdo form ^[30]^, yet it is necessary to evaluate the bovine milk extracted enzyme for its percentage of active fraction by studying the metal ion effect. However, the studies did not show any significant effect of M_O_^+6^on enzyme activity which confirms that isolated enzyme existed in the molybdo form (Figure 1a). On the contrary, EDTA inhibited the enzyme activity owing to the complex formation with M_O_^+6^at the catalytic centerconfirming the presence of MO^+6^ and its functional involvement. Further, pH optima studies revealed that the optimum pH for the maximum enzymatic reaction was determined as 7.5 (Figure 1b). Hence, all further enzymatic reactions were performed using at pH 7.5 and the kinetic data calculated from the Michaelis-Menten equation using GraphPad Prism version 8.0 (GraphPad Software 8.0, San Diego CA) was reported in Table 1 and Figure 2.

**Table 1:**
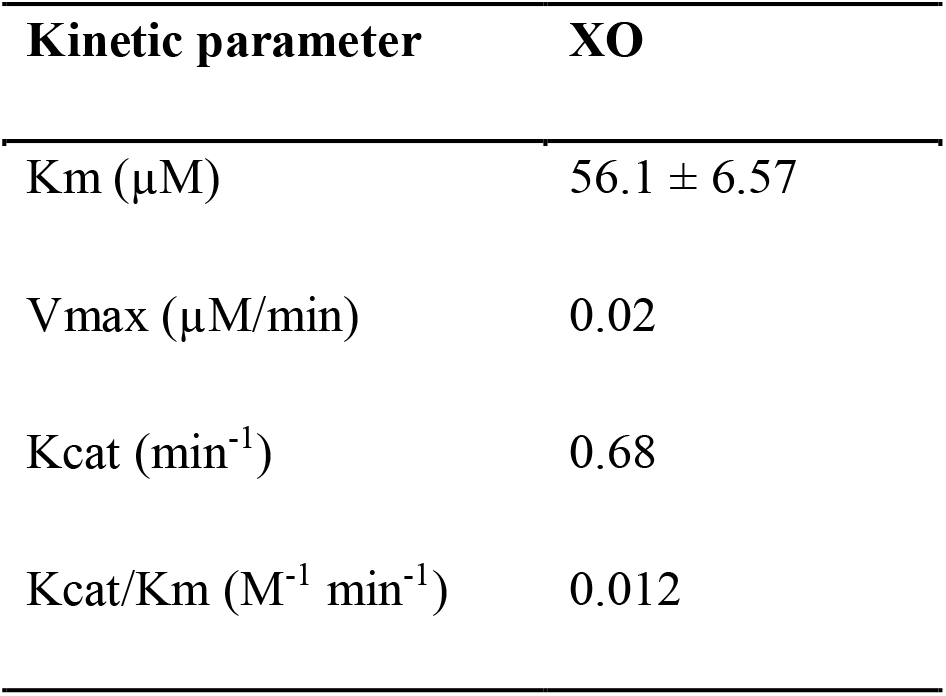
Enzyme kinetics of XO isolated from bovine milk.

| Kinetic parameter | XO |
| --- | --- |
| K<sub>m</sub> (μM) | 56.1 ± 6.57 |
| V<sub>max</sub> (μM/min) | 0.02 |
| K<sub>cat</sub> (min<sup>-1</sup>) | 0.68 |
| K<sub>cat</sub>/K<sub>m</sub> (M<sup>-1</sup> min<sup>-1</sup>) | 0.012 |

**Figure 2:**
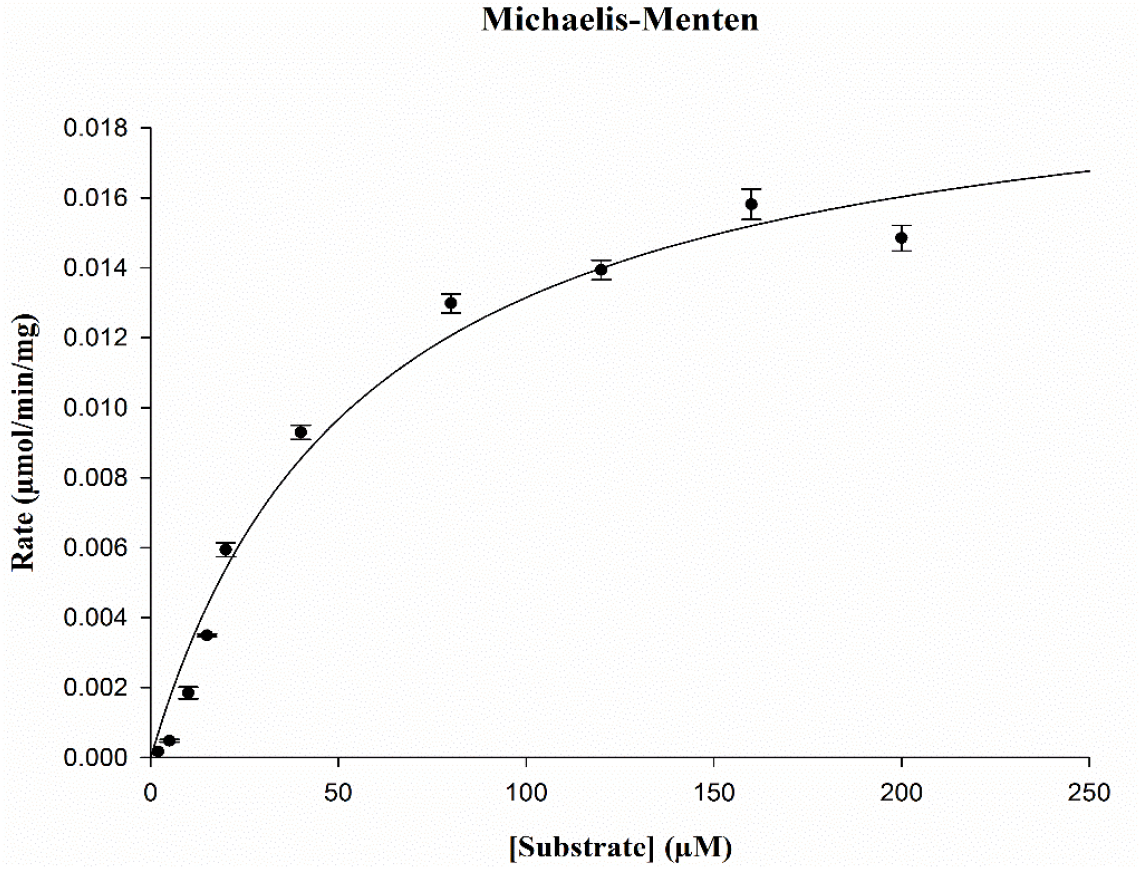
Kinetic data for xanthine against XO.

### NBT plate assay

Despite the fact that chemically synthesized molecules have been employed as a therapeutic regimen for XO inhibition, the investigation of natural compounds for their potential to XO inhibition has constantly encouraged the researchers owing to their low toxicity and potency. Profoundly, the non-purine originated XO inhibitor molecules are playing a vital role as lead molecules for future drug discovery with respect to hyperuricemia and associated disorders ^[31-32]^. Flavonoids, occupied the prominent position in the inhibition of XO ^[33]^ followed by anthraquinones (belongs to the group of flavonoids). HAQ have been classified as a multifunctional compoundswith enormous therapeutic properties ^[13]^. Though, the quinone group of compounds are well known for toxicity and mutagenicity due to generation of ROS, the intensity will be varied among the derivatives and depends on the type of the substitution. Surprisingly, Quang et al., reported the excellent superoxide scavenging potential of HAQ compared to standard quercetin and ascorbic acid which can potentiate their practice as a XO inhibitors ^[34]^. Moreover, HAQ are usually not susceptible for toxic reactions owing to their two benzene rings in α, β-positions ^[35]^, hence have advantages for their application in human health sector.

In the present study the authors screened the culture filtrate of four actinobacterial strains (*Streptomyces luteireticuli* ICTA16, *Streptomyces kurssanovii* ICTA165, *Amycolatopsis thermoflava* ICTA103and *Amycolatopsis lurida* ICTA194) for XO inhibition. Out of them, *Streptomyces luteireticuli* ICTA16, *Streptomyces kurssanovii* ICTA165, *Amycolatopsis thermoflava* ICTA103 have shown XO inhibition in the NBT plate assay. However, the results revealed the significant inhibition of XO activity (52 ± 1.27) bythe crude extract isolated from *Amycolatopsis thermoflava* ICTA 103 which is comparable to standard allopurinol (68 ± 0.5) (Figure 3 & Table 2).

**Table 2:**
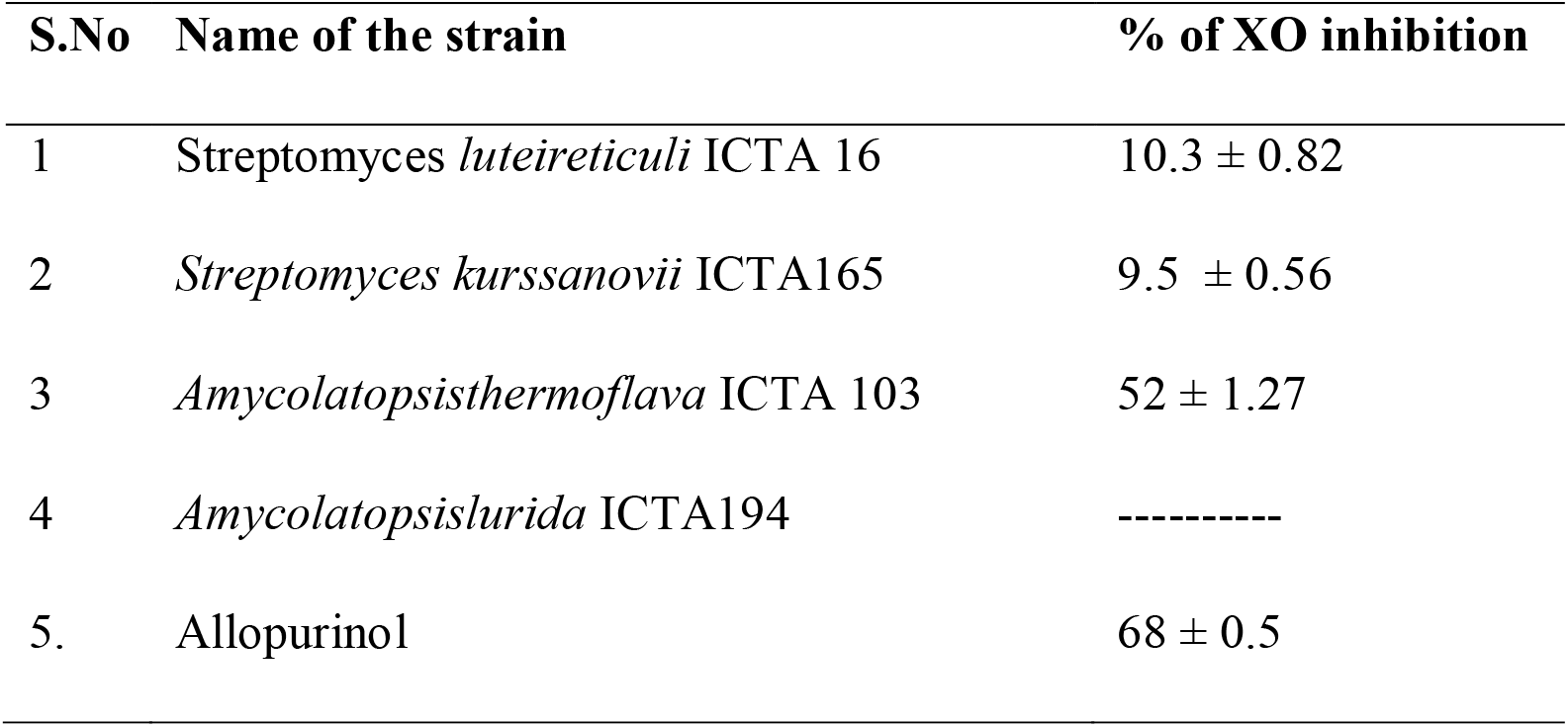
Percentage of XO inhibition of crude extracts of actinobacteria by NBT plate assay.

| S.No | Name of the strain | % of XO inhibition |
| --- | --- | --- |
| 1 | *Streptomyces luteireticuli* ICTA 16 | 10.3 ± 0.82 |
| 2 | *Streptomyces kurssanovii* ICTA165 | 9.5 ± 0.56 |
| 3 | *Amycolatopsis thermoflava* ICTA 103 | 52 ± 1.27 |
| 4 | *Amycolatopsis lurida* ICTA194 | ----- |
| 5. | Allopurinol | 68 ± 0.5 |

**Figure 3:**
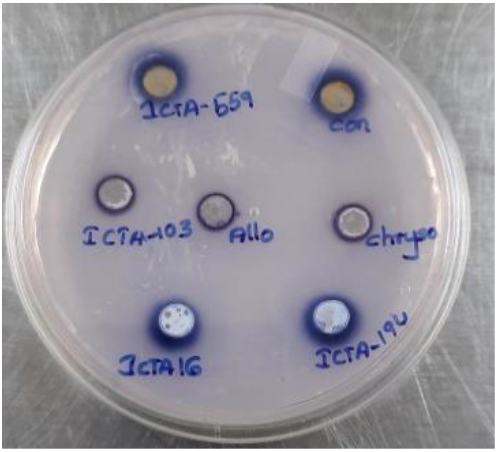
Screening of crude extracts of actinobacteriaby NBT plate assay.

### Purification and characterization of OMC from *Amycolatopsis thermoflava* ICTA 103

The crude extract of *Amycolatopsis thermoflava* ICTA 103 was used for further studies and subjected for purification of lead compound according to Kumar et al., 2017 ^[17]^. This purified compound was characterized by various analytical spectral data (NMR, FT-IR, HRMS and HPLC) and data suggested that the purified compound is similar to that reported by Kumar et al., 2017 ^[17]^, hence confirmed that the present purified XO inhibitor belongs to HAQ family and structure was identified as 1,8-Dihydroxy-3-methylanthracene-9,10-dione. Owing to its structural resemblance to chrysophanol, it is also known as 1-*O*-methyl chrysophanol(OMC).

### Molecular model building assessment

The outcomes demonstrated that OMC and chrysophanol could align similarly to quercetin in the xanthine binding pocket of the enzyme. These findings are consistent with earlier reports of mixed XO inhibitors Y700 and aloe emodin derivatives described by Shi et al., ^[36]^. The ligand’s alignment at the active pocket of the enzyme were depicted in the Figure 4.

**Figure 4:**
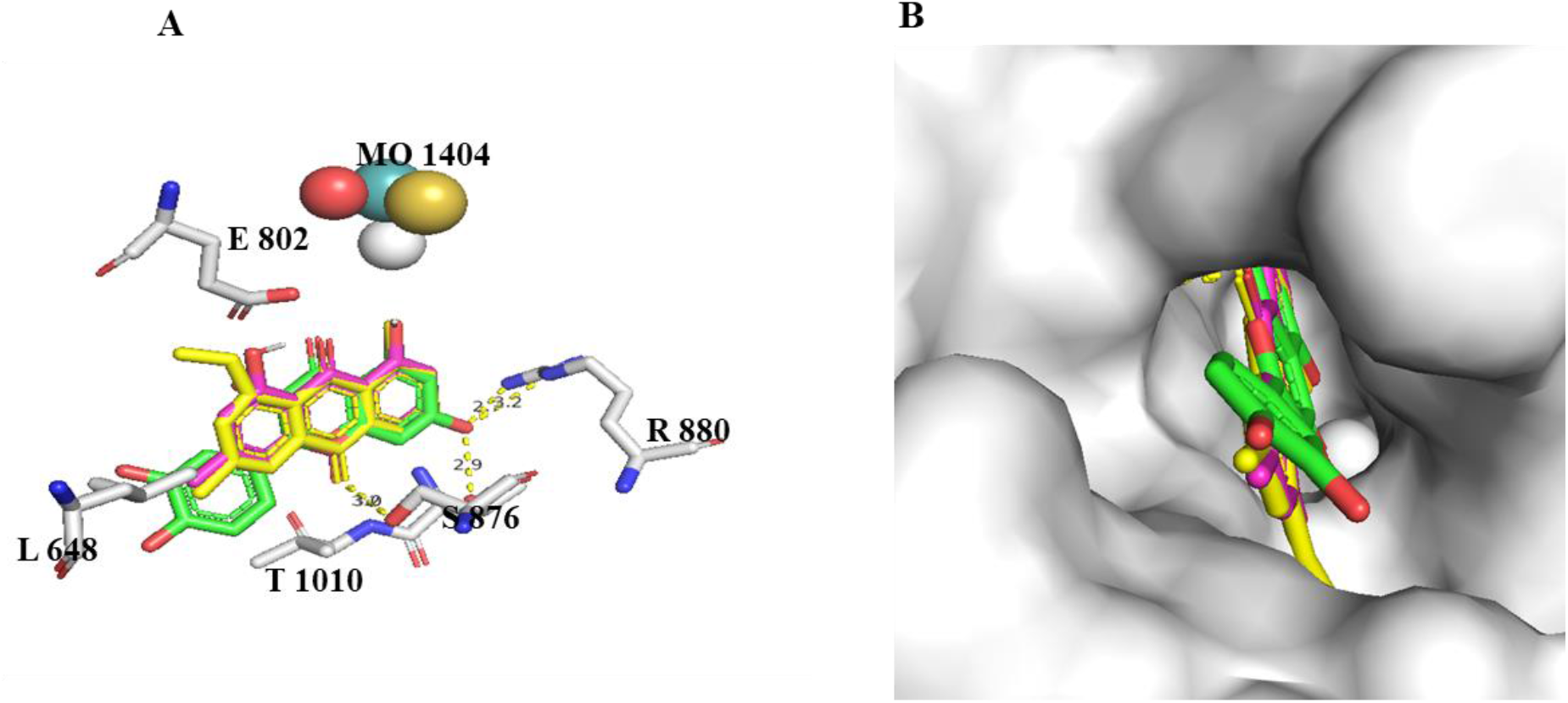
A. Closest view of potential interactions of ligands in the active pocket of XO (PDB ID: 3NVY) B. chrysophanol and OMC were docked in the binding site of XO: pink color molecule chrysophanol, green colour quercetin, yellow colour OMC (Surface model).

However, the hydrogen bonding interactions showed that quercetin effectively formed 3 hydrogen bonds with ARG 880 (2) and THR 1010 (1), whereas OMC and chrysophanol formed one hydrogen bond with SER 876. The 9-carbonyl atom of OMC and chrysophanol formed a hydrogen bond with serine 876 [figure 5]. However, the methoxy group present in the OMC exhibits hydrophobic interaction with surrounding amino acids of the active site could be the reason behind the better inhibition of OMC over chrysophanol, a structural homologue of OMC. These results mimic the previous literature of aloe-emodin and its derivatives. The crystal structure of the XO revealed that ARG 880, GLU 802, PHE 1009, PHE 914, ALA 1078 THR 1010, VAL 1011, LEU 1014 and SER 876 are the key amino acids of the active site and it is revealed from previously reported research that most of the natural inhibitors and allopurinol interacted with these residues to display their inhibition ^[37]^. Even the effective binding of 9,10-anthraquinone moiety in the outer region of XO active site is a prerequisite for the inhibition^[38]^. These findings supported the inhibition of XO by chrysophanol and OMC.

**Figure 5:**
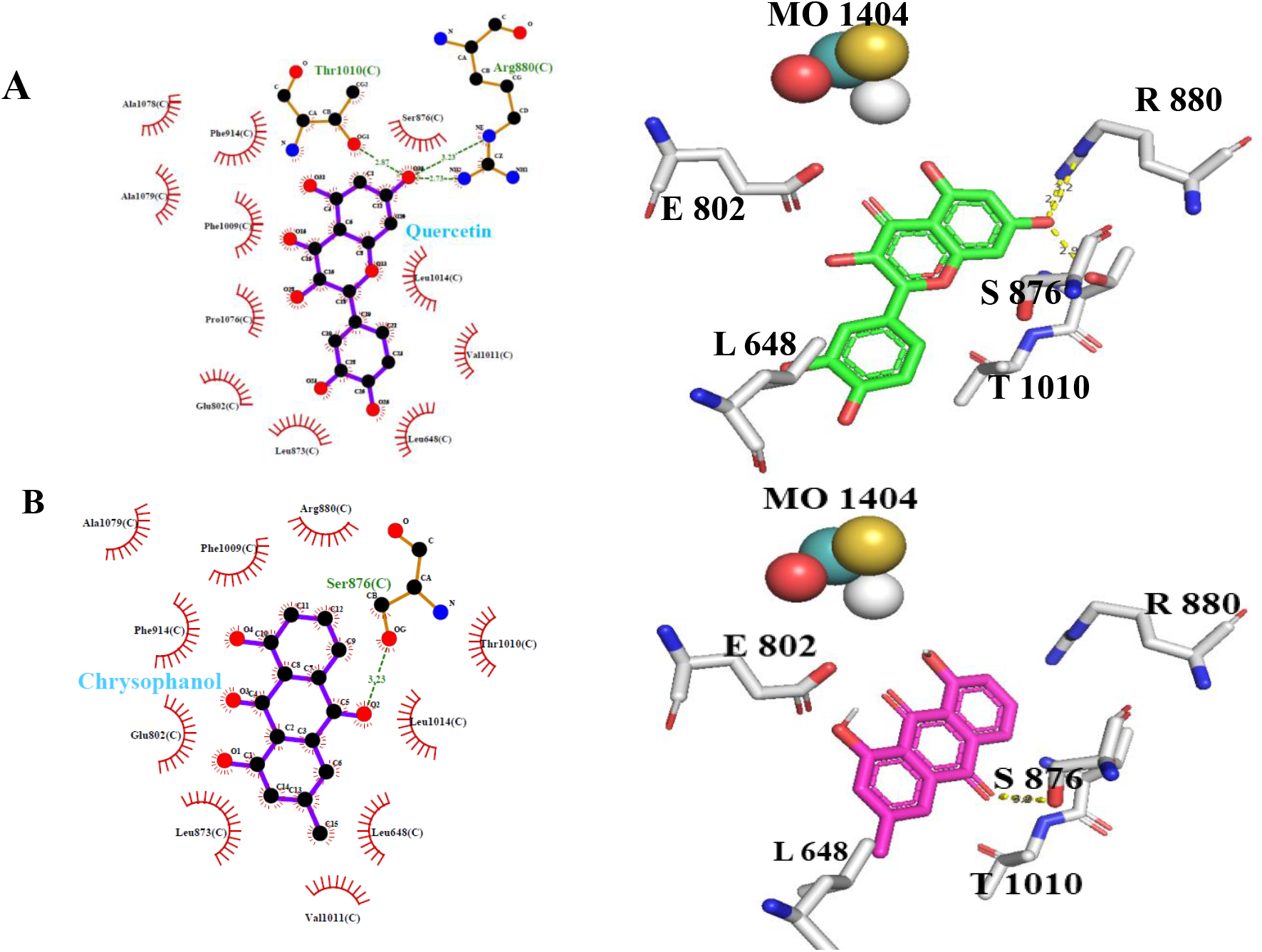
Binding pocket of the ligands in the XO. A) Quercetin B) Chrysophanol C) OMC.

### Quantitative XO inhibition studies

Kumar et al, (2017) reported that OMC is the potent superoxide radical scavenger ^[17]^ based on the non-enzymatic superoxide radical scavenging approach. However, all the superoxide scavengers may not be the XO inhibitors^[39]^.To confirm whether OMC is superoxide radical scavenger or it inhibits generation of superoxide radicals, an enzymatic assay was performed using xanthine-Xanthine oxidase system. The data revealed a significant superoxide radical scavenging (82.8 ± 0.54%). The allopurinol, as positive standard andchrysophanol as a structural standard depicted 91.7 ± 0.23 as well as 40.5 ± 1.17% of inhibition, respectively. The superoxide scavenging potential of OMC by enzymatic method was displayed in Figure 6.

**Figure 6:**
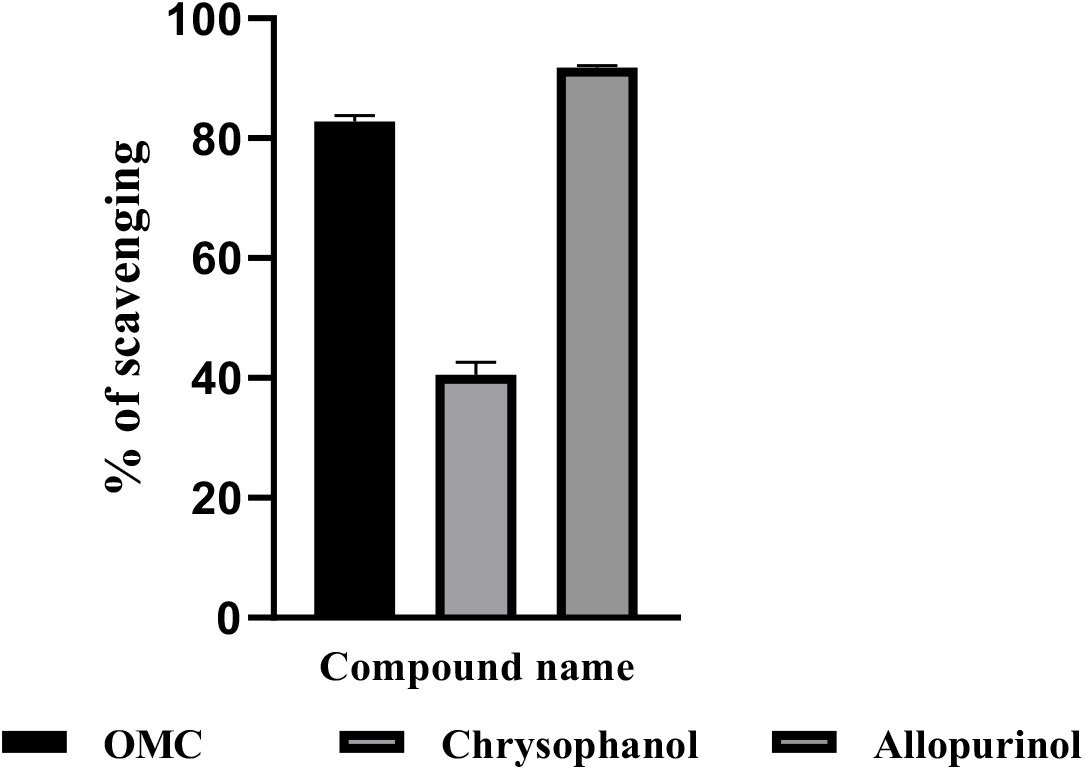
Superoxide scavenging potential of OMC using Xanthine-XO system. The results were expressed as Mean ± SD.

Critical evaluation of the above data further indicated that OMC compound inhibits the superoxide radicals at two levels; a) at the source of superoxide radical production and b) scavenging of the produced radicals. This can be attributed based on the fact that superoxide radicals are the byproducts of xanthineto uric acid conversion step catalyzed by XO and OMC being an enzyme inhibitor. This can be further confirmed with literature reports that XO is the significant producer of superoxide free radicals along with the uric acid production ^[40, 23]^. The second activity of the OMC i.e., scavenging of superoxide radicals is well documented by earlier reports ^[17]^.Thus the data on non-enzymatic scavenging assay and enzymatic scavenging assay supports that OMC could act as a two sword knife which can prevent production of superoxide radicals as well as promotes the scavenging of generated free radicals.

Thereafter, the effect of OMC on *in vitro* uric acid production was studied and the prominent results were observed. The results explored the significant percentage of XO inhibition against OMC (76 ± 0.50) which is comparable to allopurinol (98.12 ± 0.69) and higher than the chrysophanol (40.86 ± 0.65).The data was represented in the Figure 7.

**Figure 7:**
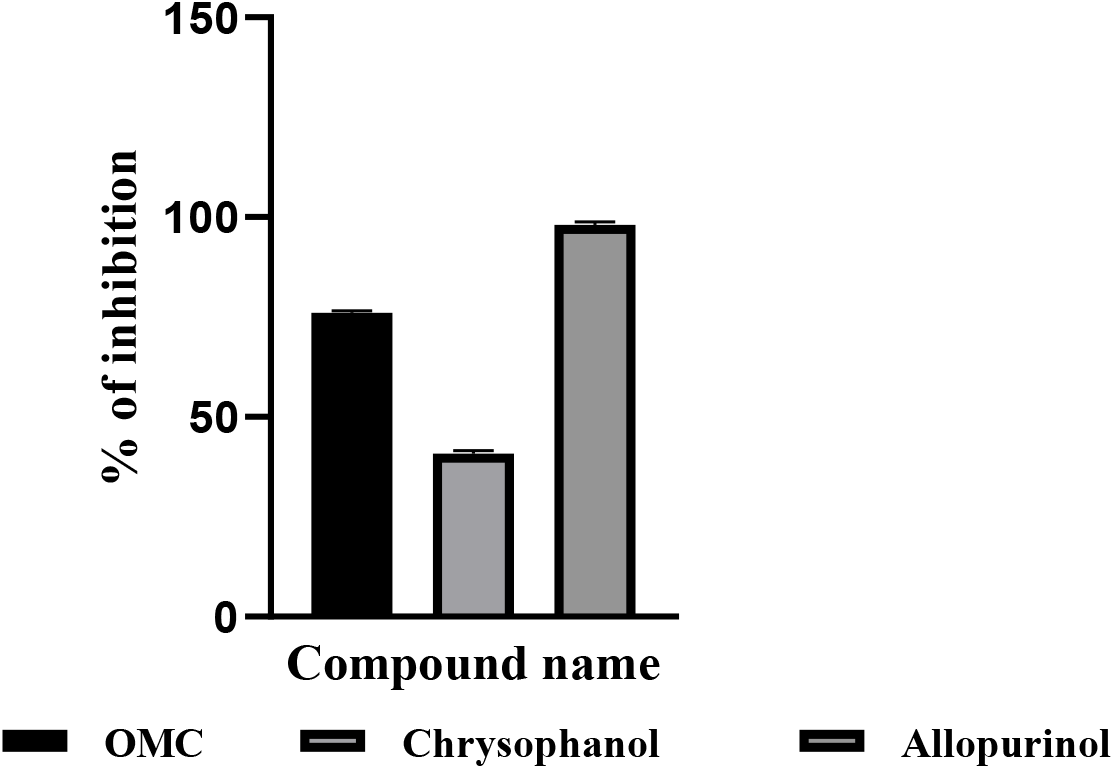
Percentage XO inhibition potential of OMC against XO.

Consecutively, the IC_50_values & inhibition constant (*K*_*i*_) values of OMC were calculated and displayed as 24.8±0.072 µM and 2.218±0.3068 µM respectively. The data was tabulated (Table 3) and graphs were represented in the figure 8& 9.

**Table 3:**
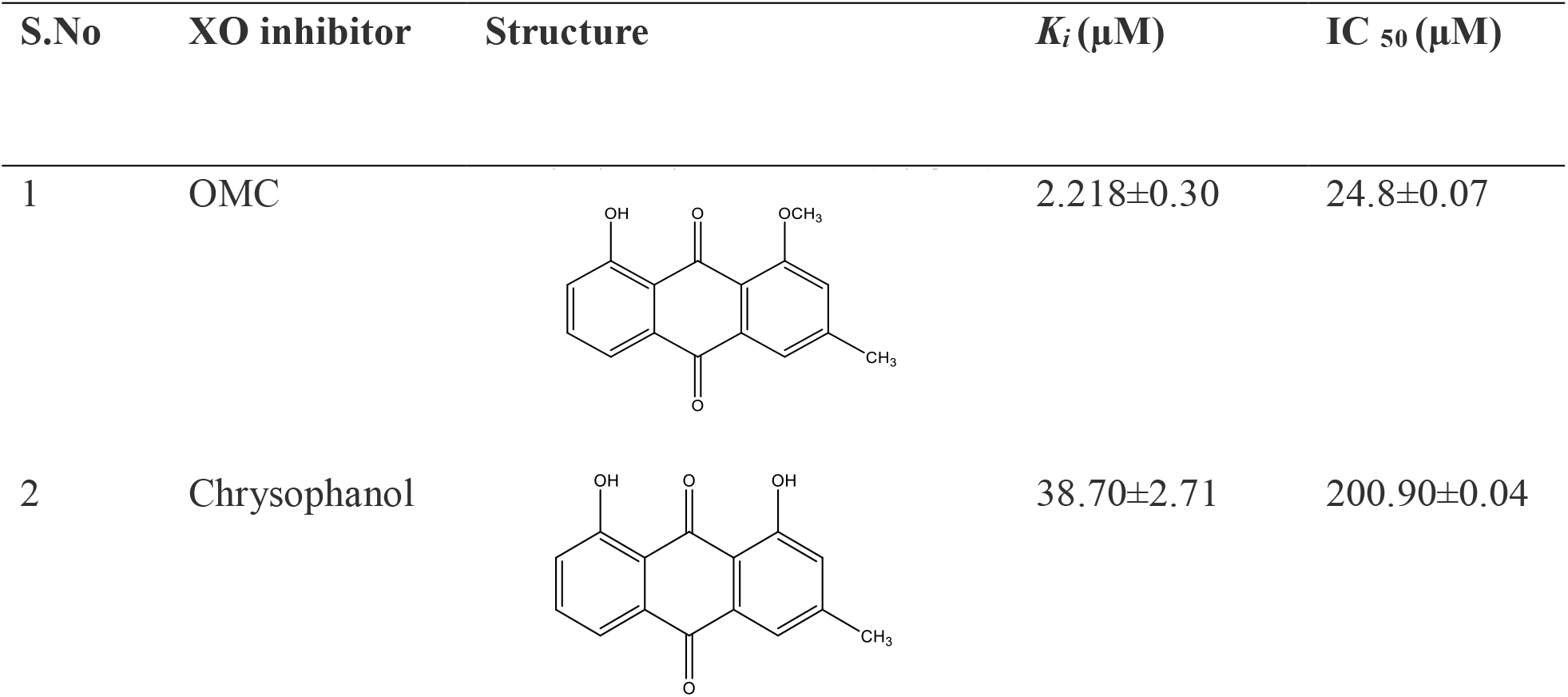
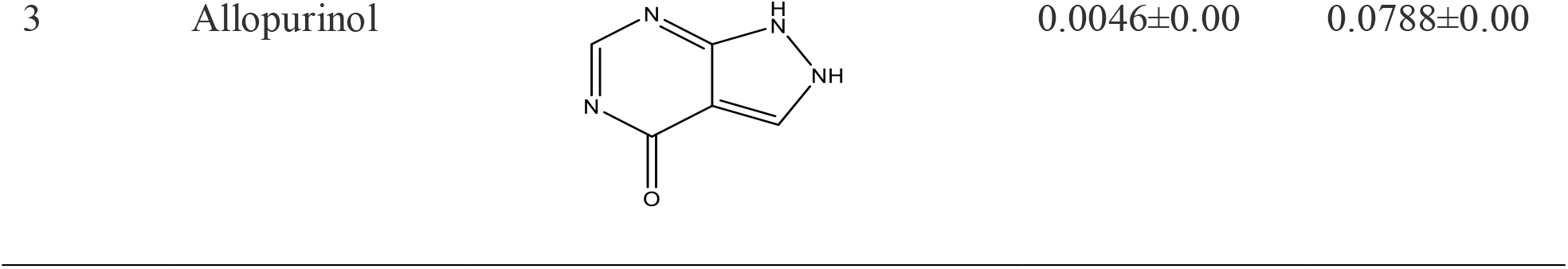
The inhibition constant *K*_i_ and IC_50_values of XO inhibitors. The Morrison *K*_i_ plots and IC_50_ plots were represented in figure 7& 8.

**Figure 8:**
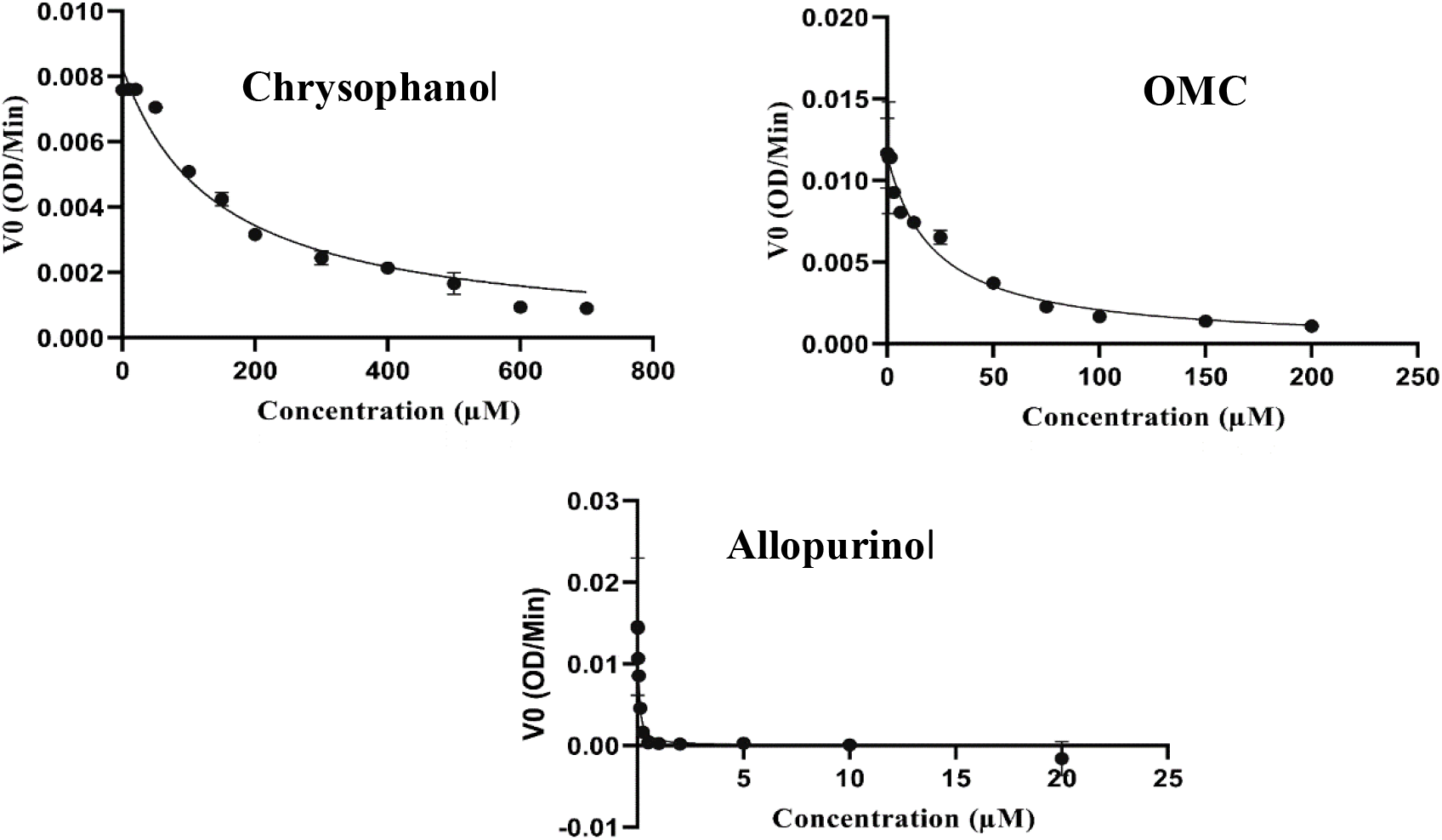
Morrison *K*_*i*_ plots for XO against inhibitors: Chrysophanol, OMC and Allopurinol.

**Figure 9:**
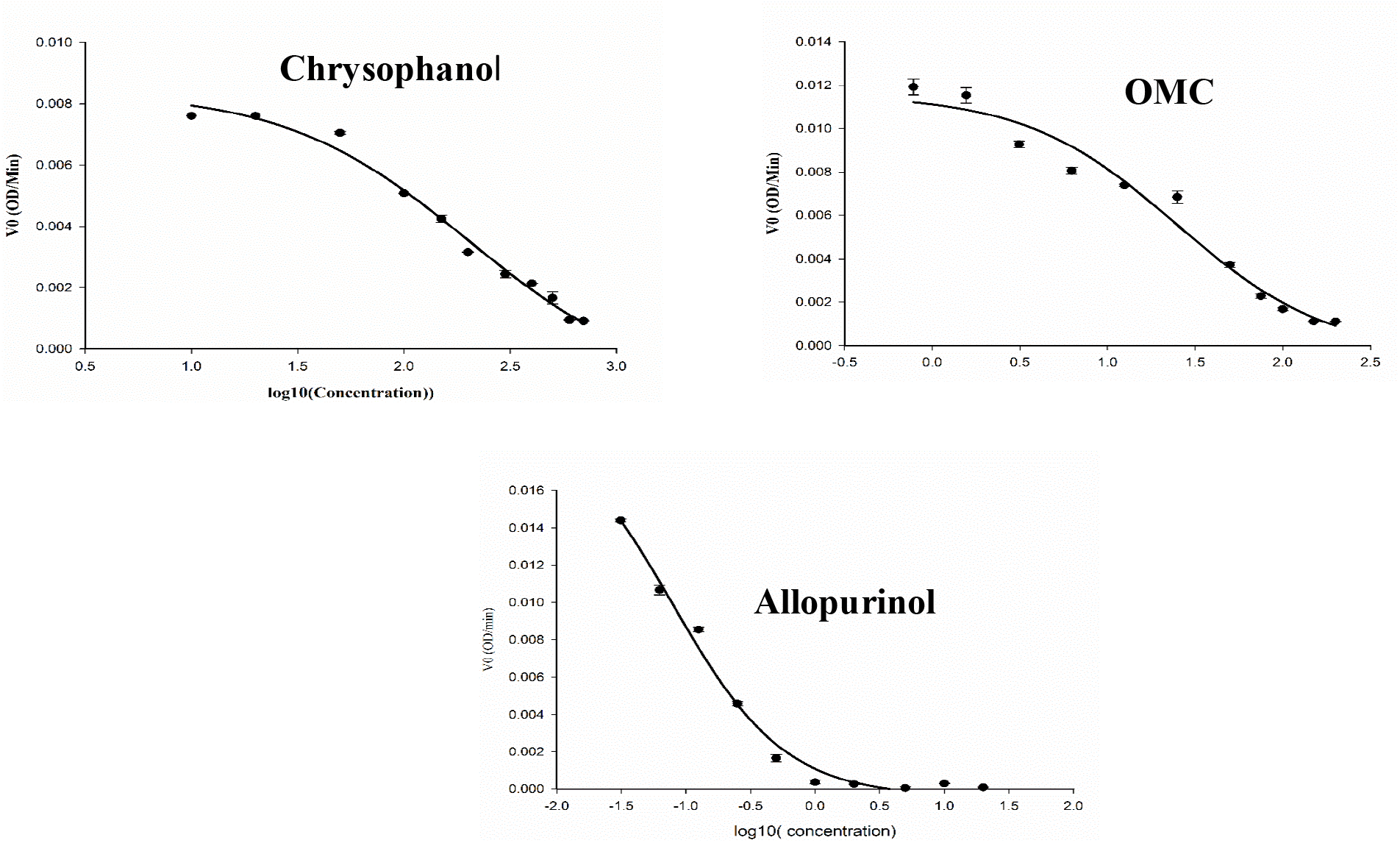
IC _50_ plots for XO against inhibitors: Chrysophanol, OMC and Allopurinol.

Former, different HAQ such as anthragallol, alizarin, rufigallol, chrysazine, aloe-emodin, emodin, physcion, chrysophanol, anthrarobin, purpurin have studied for inhibition against XO. Among them anthragallol ^[41]^, anthrarobin, purpurin ^[42]^ chrysophanol and physcion ^[43]^ had shown significant inhibition of XO with an IC_50_ values of 12, 68.35, 105.13, 143.3 and 158.5 µM respectively. Preceding, in the present study the OMC isolated from *Amycoloptosis thermoflava* ICTA 103 has shown potent XO inhibitory activity than reported HAQwith an IC_50_ value of 24.8±0.072.Eventhough, the structural difference is attributed to the presence of methoxy group in the OMC at the 1^st^ position of be anthraquinone ring, there has been significant difference in the biological activity compared to chrysophanol and other HAQ. The rise in the biological potency of the OMC may be attributed to its increased solubility in DMSO owing to its methoxy group. The theory was explained on the basis of stabilizing the development of charge separation in DMSO. The presence of a single electronegative atom (OH) in the OMC will show less charge separation which ultimately improves the DMSO solubility of the compound ^[44]^. On top of that, the reduced toxicity profile of OMC ^[14]^ compared to chrysophanol might excel in its utility as a therapeutic agent in future clinical studies.

### Mode of enzyme inhibition

In view of the above data on the efficacy and significance of OMC against XO inhibition like allopurinol and higher potency than its structural analog chrysophanol, further studies were designed to explore the mechanistic pathway by performing time dependent enzyme inhibition in presence of OMC. A pre-incubated enzyme-OMC was required for effective inhibition of enzyme by OMC and chrysophanol and lack of incubation showed minor inhibition. However, the pre-incubation did not influence the enzyme inhibition potential by allopurinol (Table 4).

**Table 4:**
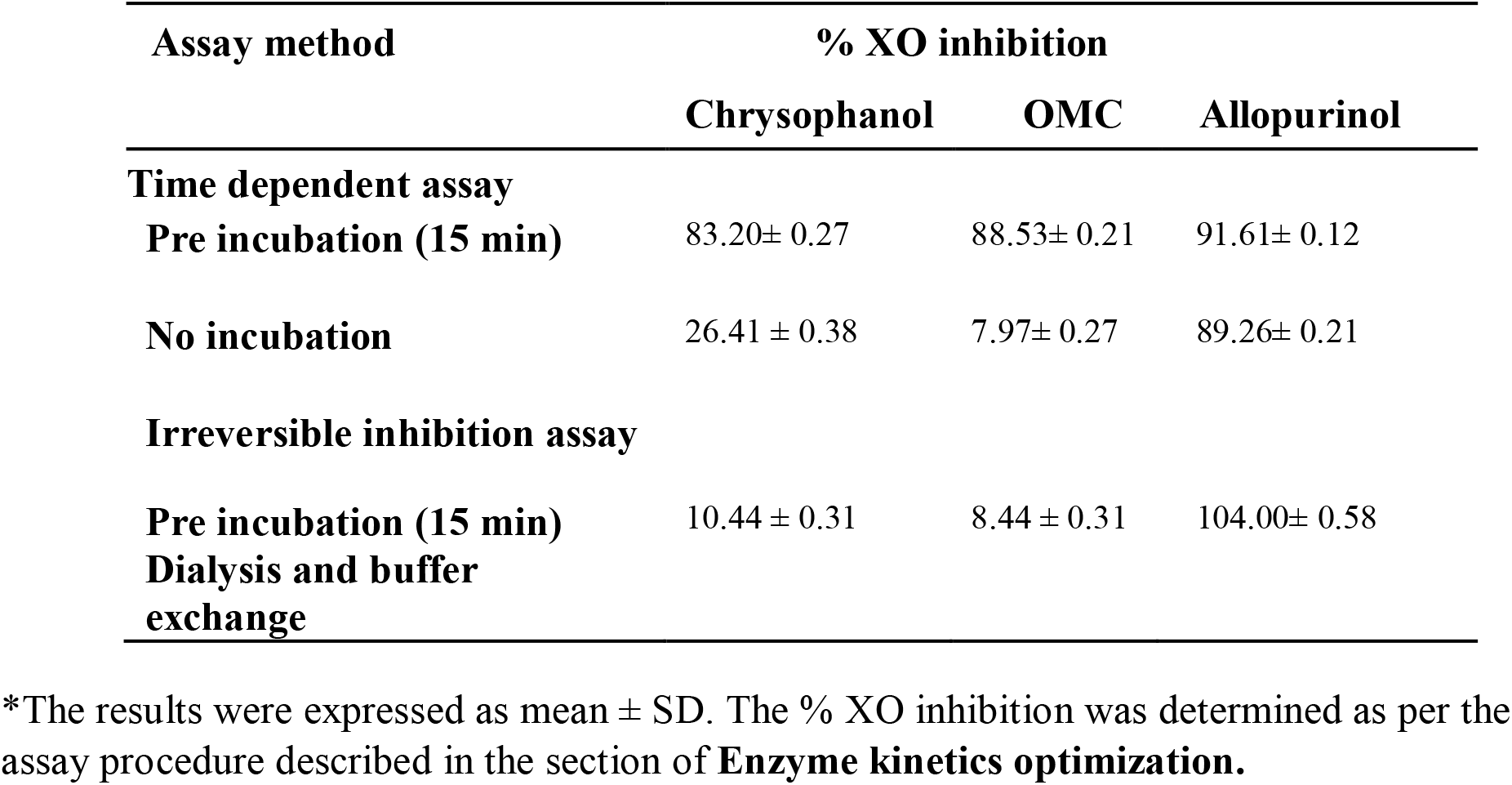
Elucidation of XO inhibition mechanism.

These results suggested the slow binding inhibition mechanism of these compounds and their ability to induce conformational changes in the enzyme to prevent the solvent accessibility to the target site. This is the first report of its kind on time dependent XO inhibition by OMC.

The reversible inhibition assay results revealed the reversibility of the potential covalent modification of XO by OMC and chrysophanol, whereas, allopurinol was fully inhibited even upon the removal of inhibitor by dialysis (Table 4). These findings displayed the reversible modification of XO by OMC and chrysophanol,however, allopurinol exhibited irreversible covalent modification by allopurinol.

### Lineweaver – Burk plots

The Lineweaver-Burk plots of OMC and chrysophanol showed an intersection at X-axis and the values of Km &Vmax were decreased as the concentration of inhibitor increased (Fig 10). This suggested the mixed type of inhibition by OMC and chrysophanol. On the contrary, the Lineweaver-Burk plot of allopurinol had intersected the X-axis displaying an increased Km value and unaltered Vmax as the inhibitor concentration increased. It proved the typical competitive inhibition of allopurinol against XO.

**Figure 10:**
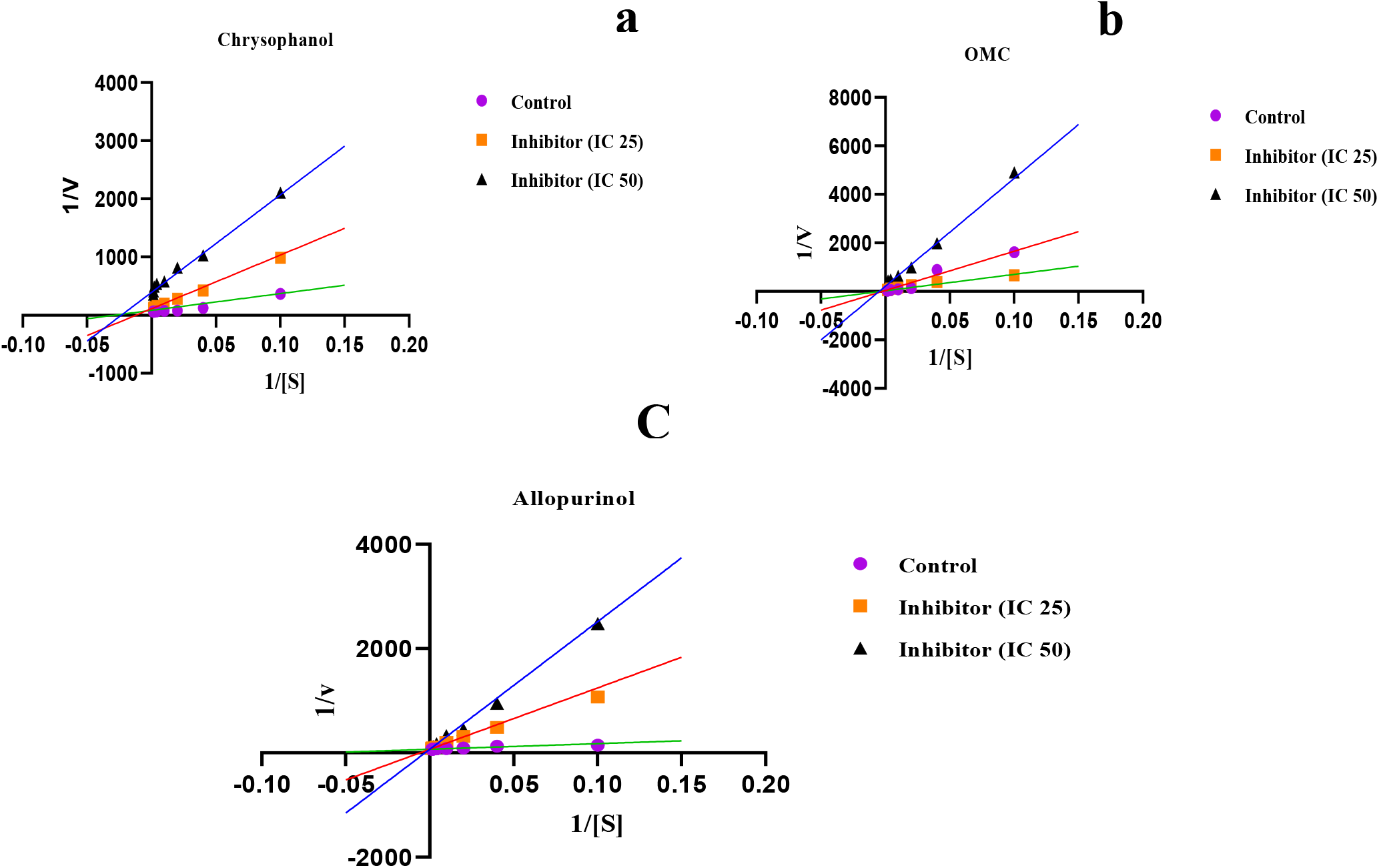
Inhibitory mechanism of inhibitors against XO enzyme. a) Chrysophanol b) OMC c) Allopurinol using Lineweaver-Burk plot (double reciprocal plot).

## Conclusion

A potent XO inhibitor was isolated from *Amycolatopsis thermoflava* ICTA103 and targeted against XO which was isolated and purified from bovine milk. The purified compound was identified as OMC based on the analytic spectral data. Simultaneously, the XO was characterized for its activity and kinetics using xanthine as substrate and noticed that it has Km 56.1 ± 6.57 µM, Vmax 0.020 µM/min and Kcat 0.68 min^-1^. Further, *in silico*molecular model building studies indicatedthe significant binding interaction between XO and OMC in the active pocket which is further confirmed by *in vitro*enzyme inhibition studies. The *in vitro* XO inhibition studies explored the significant % inhibition of OMC (76±0.61) which is comparable to standard allopurinol (98.12 ± 0.69). Moreover, the significant IC_50_ (24.8 ± 0.072 µM) & *K*_*i*_(2.218 ± 0.3068 µM) values identified it as a potential candidate for XO inhibition. The mechanism of inhibition was explored by kinetic and Lineweaver-Burkplots and recognized as reversible covalent mode of inhibition showing mixed inhibition model. Further *in vivo* XO inhibition potential of OMC using animal models and *in vitro* inhibition on cell line models are in progress.

## Acknowledgments

Authors are thankful toDr. Sunil Misra, Dr. Kiranmai Mandava, Dr. Anthony Addlagatta and Dr. Ramars Amanchy for their constant support and encouragement and CSIR-New Delhi for funding under the Emeritus scheme, grant number 21(1102)/20/EMR-II.

