## Supplementary information for "Inhibition of Xanthine oxidase by 1-*O*-methyl chrysophanol, a hydroxyanthraquinone isolated from *Amycolatopsis thermoflava* ICTA 103"

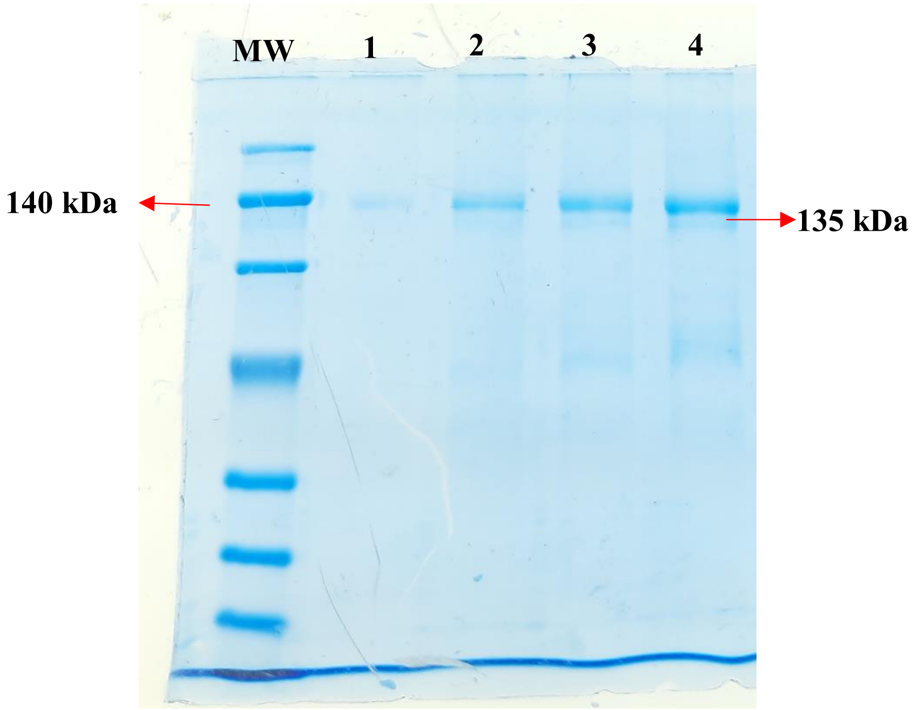


**Fig S1: SDS-PAGE of XO isolated from Bovine milk on 12% gel. Different volumes of protein loading (16mg/ml). Lane 1-1 µl, Lane 2- 2 µl, Lane 3- 3µl, Lane 4-4 µl.**

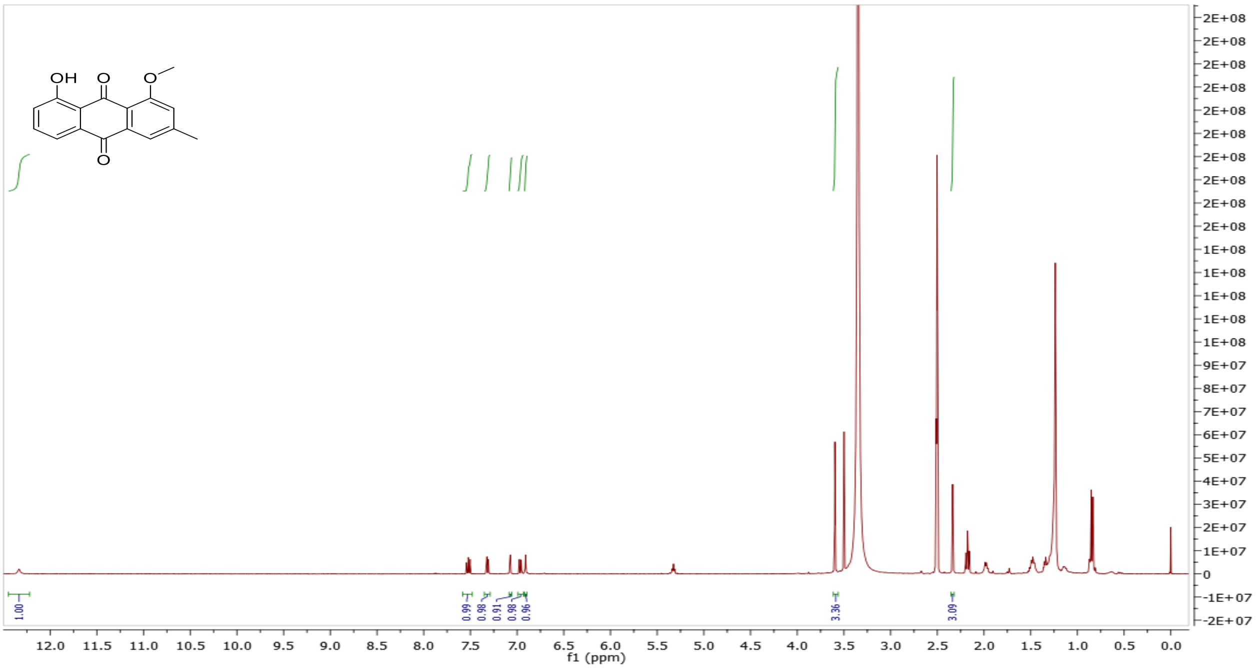

**Figure S2: $^1$H NMR spectra of OMC in DMSO-d6**

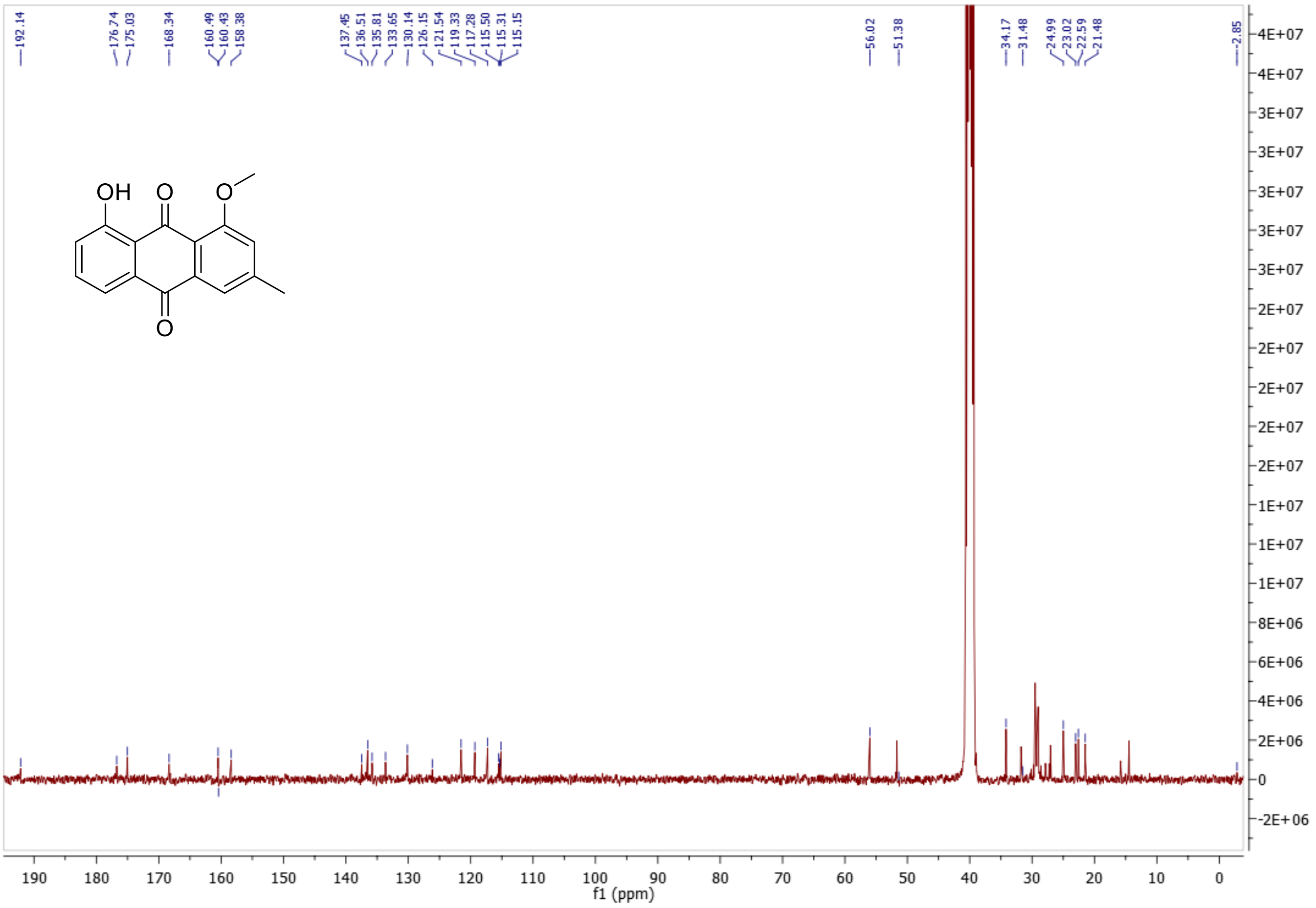


**Figure S3: $^{13}$C NMR spectrum of OMC in DMSO-d6**

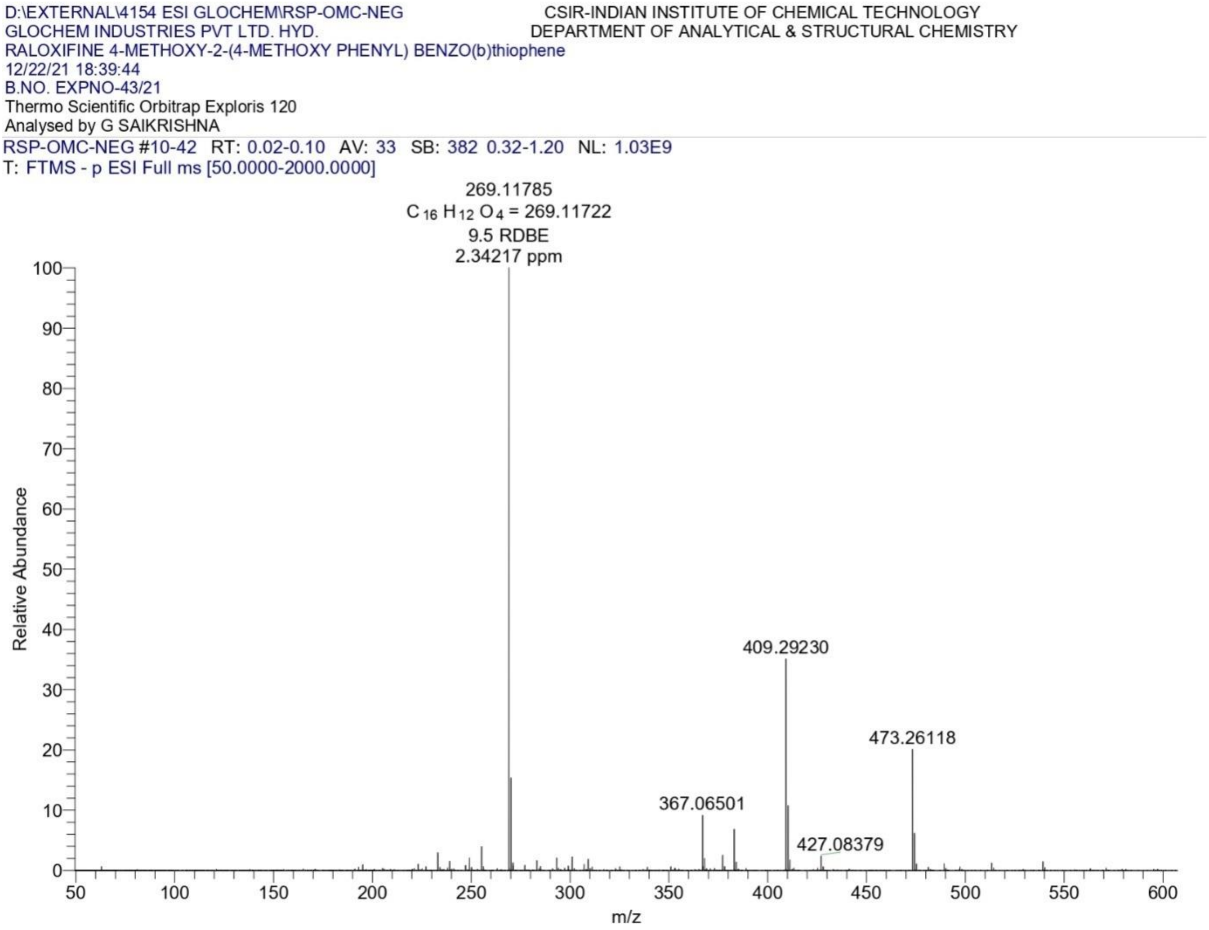


**Figure S4: HR-MS spectrum of OMC in methanol.**

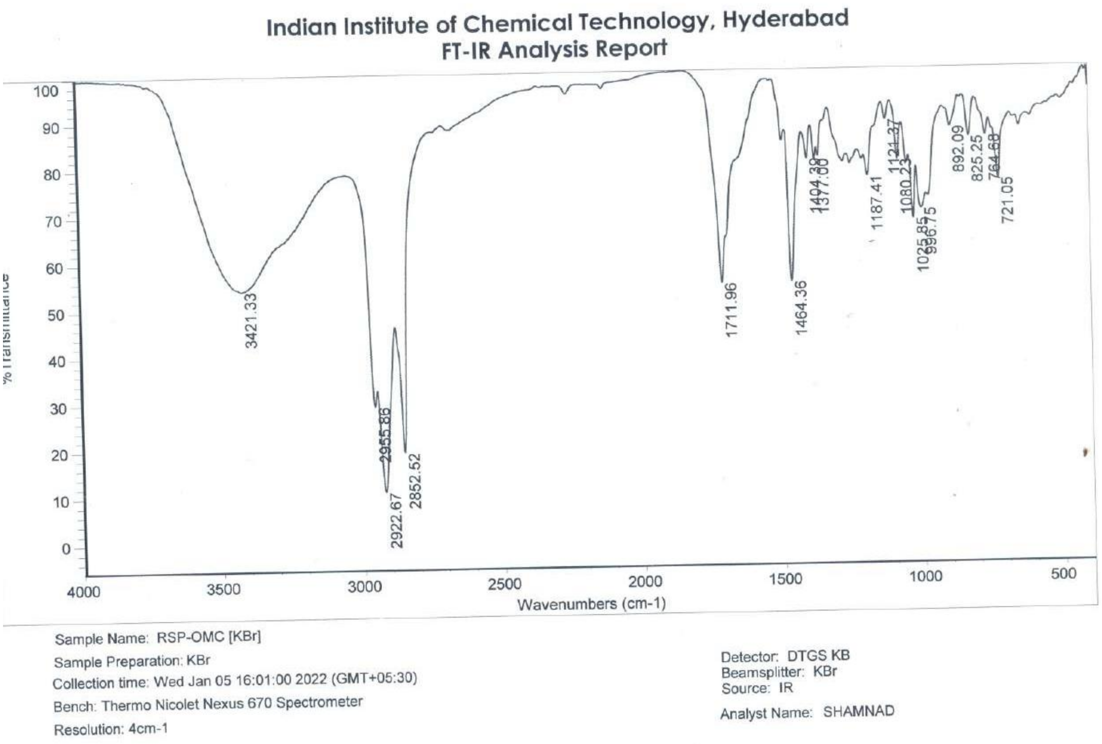


**Figure S5: FT-IR spectrum of OMC in KBr pellet.**

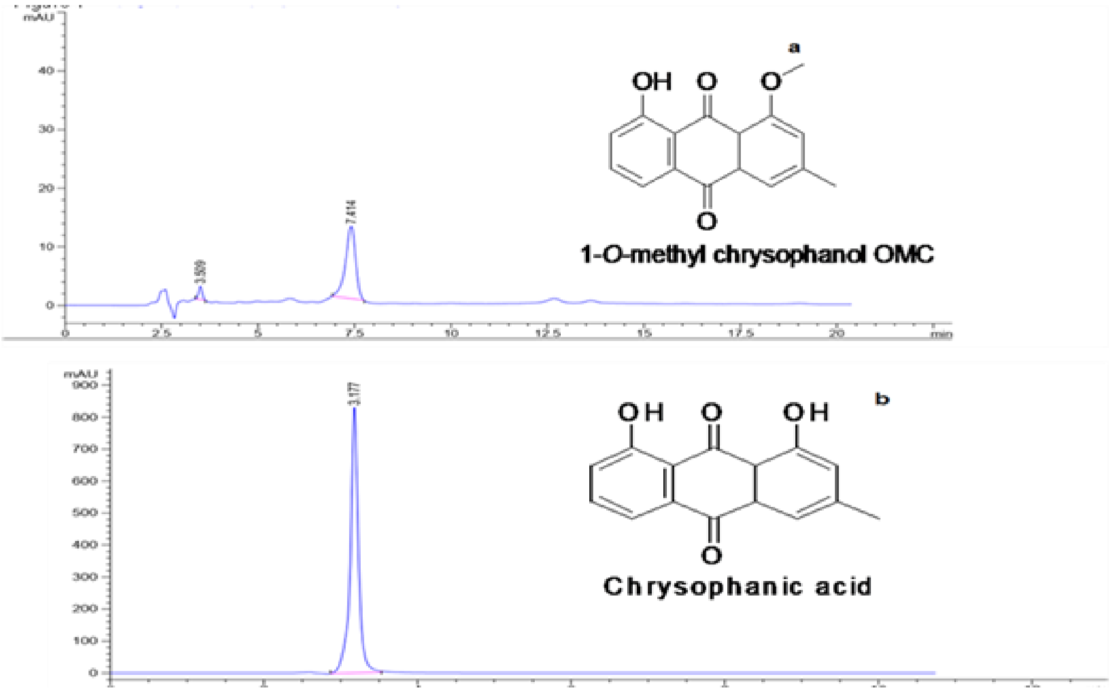


**Figure S6: HPLC chromatogram of purified OMC; Chyrsophanol (reference compound).**
